# The *Sorghum bicolor* reference genome: improved assembly and annotations, a transcriptome atlas, and signatures of genome organization

**DOI:** 10.1101/110593

**Authors:** Ryan F. McCormick, Sandra K. Truong, Avinash Sreedasyam, Jerry Jenkins, Shengqiang Shu, David Sims, Megan Kennedy, Mojgan Amirebrahimi, Brock Weers, Brian McKinley, Ashley Mattison, Daryl Morishige, Jane Grimwood, Jeremy Schmutz, John Mullet

## Abstract

*Sorghum bicolor* is a drought tolerant C4 grass used for production of grain, forage, sugar, and lignocellulosic biomass and a genetic model for C4 grasses due to its relatively small genome (~800 Mbp), diploid genetics, diverse germplasm, and colinearity with other C4 grass genomes. In this study, deep sequencing, genetic linkage analysis, and transcriptome data were used to produce and annotate a high quality reference genome sequence. Reference genome sequence order was improved, 29.6 Mbp of additional sequence was incorporated, the number of genes annotated increased 24% to 34,211, average gene length and N50 increased, and error frequency was reduced 10-fold to 1 per 100 kbp. Sub-telomeric repeats with characteristics of Tandem Repeats In Miniature (TRIM) elements were identified at the termini of most chromosomes. Nucleosome occupancy predictions identified nucleosomes positioned immediately downstream of transcription start sites and at different densities across chromosomes. Alignment of the reference genome sequence to 56 resequenced genomes from diverse sorghum genotypes identified ~7.4M SNPs and 1.8M indels. Large scale variant features in euchromatin were identified with periodicities of ~25 kbp. An RNA transcriptome atlas of gene expression was constructed from 47 samples derived from growing and developed tissues of the major plant organs (roots, leaves, stems, panicles, seed) collected during the juvenile, vegetative and reproductive phases. Analysis of the transcriptome data indicated that tissue type and protein kinase expression had large influences on transcriptional profile clustering. The updated assembly, annotation, and transcriptome data represent a resource for C4 grass research and crop improvement.

## 3 INTRODUCTION

*Sorghum bicolor*, the fifth most important cereal crop in the world, is an economically important C4 grass grown for the production of grain, forage, sugar/syrup, brewing, and lignocellulosic biomass production for bioenergy. Meeting the food and fuel production challenges of the coming century will require production gains from traditional crop breeding, genomic selection, genome editing, and biotechnology approaches that develop plants with increased productivity and traits such as drought, pest and disease resistance, and canopies that have high photosynthetic efficiencies (Kromdijk et al., 2016; Mickelbart et al., 2015; Mondal et al., 2016; Mullet et al., 2014; Ort et al., 2015; Park et al., 2015; Technow et al., 2015; Voytas, 2013). Progress towards the genetic improvement of plants is promoted by the availability of foundational genetic and genomic resources. Because of this, we improved the *Sorghum bicolor* reference genome sequence assembly using targeted approaches and improved its annotation using data from a deep transcriptome analysis. A sorghum transcriptome atlas was created that contains gene expression data from the major plant tissue types across the juvenile, vegetative and reproductive stages of development. The genome sequence was used to analyze the distribution of key features in the genome including genes, transposable elements, genetic variation, and nucleosome occupancy likelihoods.

Sorghum is a diploid C4 grass with 10 chromosomes and an ~800 Mbp genome (Price et al., 2005). Cytogenetic and genetic analyses showed that sorghum chromosomes are comprised of distal regions of high gene density that exhibit high rates of recombination and large heterochromatic pericentromeric regions characterized by low gene density and low rates of recombination (Kim et al., 2005). A *Sorghum bicolor* reference genome sequence was reported in 2009, representing a major landmark in C4 grass genomics (Paterson et al., 2009). Reduced sequencing costs and technological advances have since enabled the sequencing and assembly of additional grass genomes, including *Brachypodium distachyon* (Vogel et al., 2010), corn (Schnable et al., 2009), foxtail millet (Bennetzen et al., 2012; Zhang et al., 2012), wheat (Brenchley et al., 2012), barley (Consortium, 2012b), and the desiccation tolerant *Oropetium thomaeum* (VanBuren et al., 2015). In addition, the genomes of 49 additional sorghum genotypes have been sequenced and assembled through alignment to the sorghum reference genome produced in 2009 (Evans et al., 2013; Mace et al., 2013; Zheng et al., 2011). Reference genomes provide an important resource for analyses, but their coverage and quality are often limited by the resources and technology available at the time of their construction. As such, reference genomes and their annotations benefit from iterative improvement as exemplified by the Human genome project and related projects such as ENCODE (Consortium, 2012a; Consortium, 2004; Lander et al., 2001; Rosenbloom et al., 2013). To this end, we report an update to the BTx623 sorghum reference genome that leverages advances in sequencing technologies and transcriptomics to generate a more complete sorghum genome assembly and annotation.

A sorghum transcriptome atlas containing expression profiles of the major plant tissues was constructed to facilitate annotation of genes in the sorghum genome. Such atlas projects serve as resources for gene discovery, annotation, and functional characterization. Multiple atlas projects have been executed in recent years, including for maize and rice (Sekhon et al., 2013; Sekhon et al., 2011; Wang et al., 2010). In sorghum, microarray-based expression profiling and RNAseq have also been used to examine transcriptome dynamics in different sorghum genotypes, tissues, and responses to hormones and the environment (Abdel-Ghany et al., 2016; Shakoor et al., 2014). The current study contributes additional information on sorghum gene expression through construction of a sorghum transcriptome atlas using 47 samples collected from the major plant tissue types during the juvenile, vegetative and reproductive phases of plant development. Here we utilize the sorghum transcriptome atlas to facilitate gene annotation and to identify genes important for establishing organ identity in sorghum.

Additional features of the sorghum genome were investigated, including repetitive DNA elements, primary sequence-based nucleosome occupancy likelihoods, and the distribution of genetic variation among diverse sorghum accessions. Of particular interest was the identification of signatures that reflect higher-level organizational properties of the genome. Genetic variants do not accumulate uniformly across the genome due in part to regional variation in mutation rates (RViMR) that over time cause large differences in the number of genetic variants in different regions of eukaryotic genomes (Evans et al., 2013; Hodgkinson and Eyre-Walker, 2011; Makova and Hardison, 2015; Tolstorukov et al., 2011). In particular, chromatin structure has been associated with variation in the accumulation of genetic variants in human genomes (Tolstorukov et al., 2011). Additionally, previous work in medaka and humans found that genetic variation accumulated with a periodicity corresponding to nucleosome occupancy at transcription start sites (Higasa and Hayashi, 2006; Sasaki et al., 2009). Since nucleosome occupancy is associated with sequence identity, a support vector machine (SVM) was previously trained on human chromatin to predict nucleosome occupancy likelihoods from primary sequence, and the same SVM was shown to perform well in maize in predicting nucleosome occupancy (Fincher et al., 2013; Gupta et al., 2008). Given that eukaryotic genomes are organized into higher order topologically associating domains and the influence of nucleosome occupancy on the accumulation of genetic variation, the possibility that larger chromatin domains influence the genome in a similar manner in plants also exists (Bonev and Cavalli, 2016). As such, we explored the basis of genetic variation accumulation in the sorghum genome using digital signal processing techniques.

## 4 RESULTS

### 4.1 Genome assembly and improvement

Version 1 of the sorghum BTx623 reference genome assembly incorporated 625.6 Mbp of genomic sequence into 10 pseudomolecules corresponding to the 10 sorghum chromosomes by combining data from whole genome shotgun sequencing and targeted sequencing of BACs and fosmids using paired-end Sanger sequencing,. An error rate of < 1 per 10 kbp was estimated based on Sanger sequencing of BACs (Paterson et al., 2009). Version 2 of the sorghum reference genome assembly was publicly released without a corresponding publication; as such, all comparisons here are made relative to version 1.

In this study, version 1 of the sorghum reference genome was refined by deep whole genome short read sequencing (110X) and targeted finishing of gene-dense regions of the genome (greater than 2 genes per 100 kbp) using primer walking via Sanger sequencing and shotgun sequencing of plasmid subclones, fosmid, and BAC clones (Supplemental File S1). These finished regions were assembled and hand-curated (representing 344.4 Mbp), mapped back to the v1 assembly, and then incorporated into the v1 assembly, adding a total of 4.96 Mbp to the assembly. To improve ordering of the reference genome, a high-density genetic map based on ~10,000 markers genotyped in a 437-line recombinant inbred mapping population derived from the sorghum lines BTx623 and IS3620C was used to integrate 7 additional scaffolds into chromosomes (Truong et al., 2014). Furthermore, the genetic map identified a 1.08 Mbp region that was previously assembled into chromosome 6, but markers within the region were not linked to flanking regions on chromosome 6 and tightly linked with markers on chromosome 7 (Figure 1). This assembly error in version 1 is corrected in version 3.

**Figure 1:**
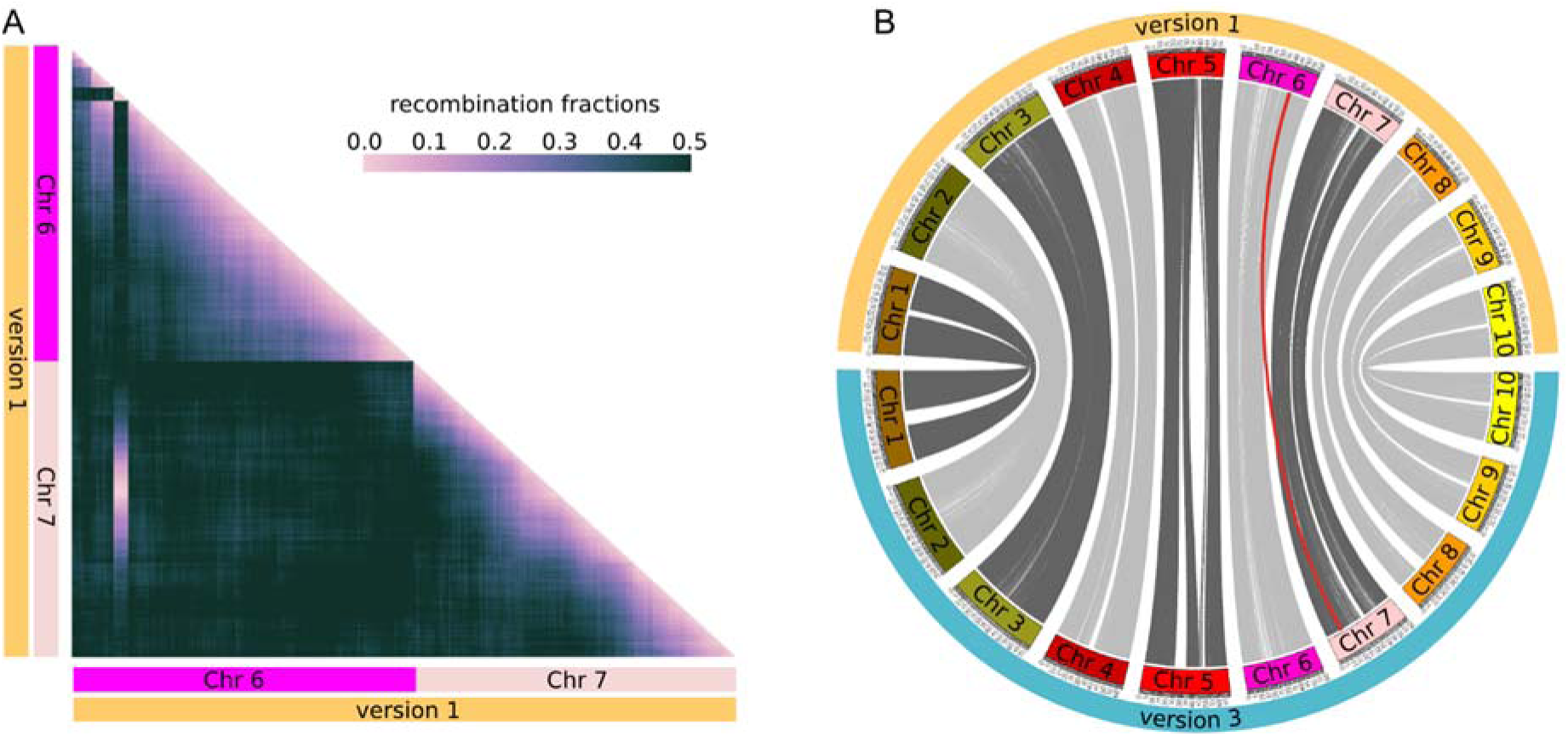
Correction of misassembled region in the version 1 sorghum reference genome assembly and integration of new sequence. (A) Recombination fractions of markers in the BTx623 x IS3620C sorghum recombinant inbred line (RIL) population ordered by physical position relative to the version 1 reference assembly. A block of markers spanning roughly 1 Mbp were previously physically assembled on chromosome 6, but are genetically unlinked with markers on chromosome 6. Instead, the markers are tightly linked with a region of chromosome 7. (B) Sequence identity mapped between the version 1 and version 3 of the reference assemblies. A 1.08 Mbp region previously located on chromosome 6, corresponding to the markers in panel A, was moved to chromosome 7. Additional sequences were integrated into the chromosomes, expanding the size of the version 3 assembly (Supplemental File S1).

Due to integration of additional sequence during finishing and of previously unplaced contigs into the main genome sequence, the contiguity of the v3 sequence comprising the 10 sorghum chromosomes increased significantly, such that the N50 length, the largest length such that 50% of all bases are contained in contigs of at least that length (Lander et al., 2001), increased by 6.3 fold from 0.2045 Mbp to 1.5 Mbp. The resulting v3 assembly included 655.2 Mbp of genomic sequence incorporated into chromosomes, with an estimated error rate of <1 per 100 kbp (Table 1).

**Table 1:** Summary statistics for sequence comprising the 10 chromosomes for the version 1 and version 3 reference assemblies. The number of bases incorporated into the genome, the contiguity of the sequence, and the accuracy of the sequence improved in version 3. N50 is defined as the largest length such that 50% of all bases are contained in contigs of at least that length (Lander et al., 2001), and L50 is defined as the number of contigs, where, when summed longest to shortest, the sum exceeds 50% of the assembly size.

|  | *Sorghum bicolor* reference genome pseudomolecules |  |
| --- | --- | --- |
|  | Version 1 | Version 3 |
| Number of pseudomolecules | 10 | 10 |
| Number of contigs | 6,929 | 2,688 |
| Scaffold sequence (Mbp) | 659.2 | 683.6 |
| Contig sequence (Mbp) | 625.6 | 655.2 |
| Scaffold N50 (Mbp) | 64.3 | 68.7 |
| Contig N50 (Mbp) | 0.2045 | 1.5 |
| Scaffold L50 | 5 | 5 |
| Contig L50 | 838 | 71 |
| Unmapped sequence (Mbp) | 71.9 | 20.2 |
| Estimated error rate | < 1 per 10 kbp | < 1 per 100 kbp |

### 4.2 Annotation of genes and other features in the sorghum genome

The version 3 (v3.1) assembly was annotated for a number of feature types, including genes, repetitive elements, genetic variation, and primary sequence-based nucleosome occupancy predictions (Figure 2, Supplemental Figures S1 and S2). Deep transcriptome profiles were obtained from 47 different tissues or developmental phases to facilitate the annotation of genes in the sorghum genome. Tissues from growing and developed portions of roots, leaves, stems, seeds, and panicles were isolated during the juvenile, vegetative, and reproductive phases of plant development. Illumina sequencing of cDNA obtained from these tissue samples (RNA-seq) generated 3.3 billion sorghum paired-end reads. The sequence reads were subsequently combined with sorghum ESTs and homology-based predictions to annotate 34,211 genes in the *Sorghum bicolor* genome (gene set version 3.1). The v3.1 gene annotation represents a 24% increase relative to the 27,607 genes annotated in version 1 (gene set version 1.4). The median and mean gene size in v3.1 increased to 1600 and 1835, from 1336 and 1473 in v1.4, respectively, due primarily to improved annotation of exons. As such, the number of genes, as well as the length of genes increased significantly indicating that the v3.1 gene annotation is the most comprehensive sorghum gene annotation to date. A small number (175) of genes in v1.4 were not supported and were not included in the v3.1 gene set. Repetitive elements in the sorghum genome were annotated using a *de novo* repetitive element annotation pipeline in conjunction with existing repetitive element libraries (Bao et al., 2015; Flutre et al., 2011; Ouyang and Buell, 2004; Quesneville et al., 2005). Consistent with the previous annotation of the v1 assembly, the percentage of the genome annotated as retrotransposons (i.e. class I elements) was 58.8%, most of which were long terminal repeats (54% of the genome). Approximately 8.7% of the genome annotated as DNA transposons (i.e. class II elements).

**Figure 2:**
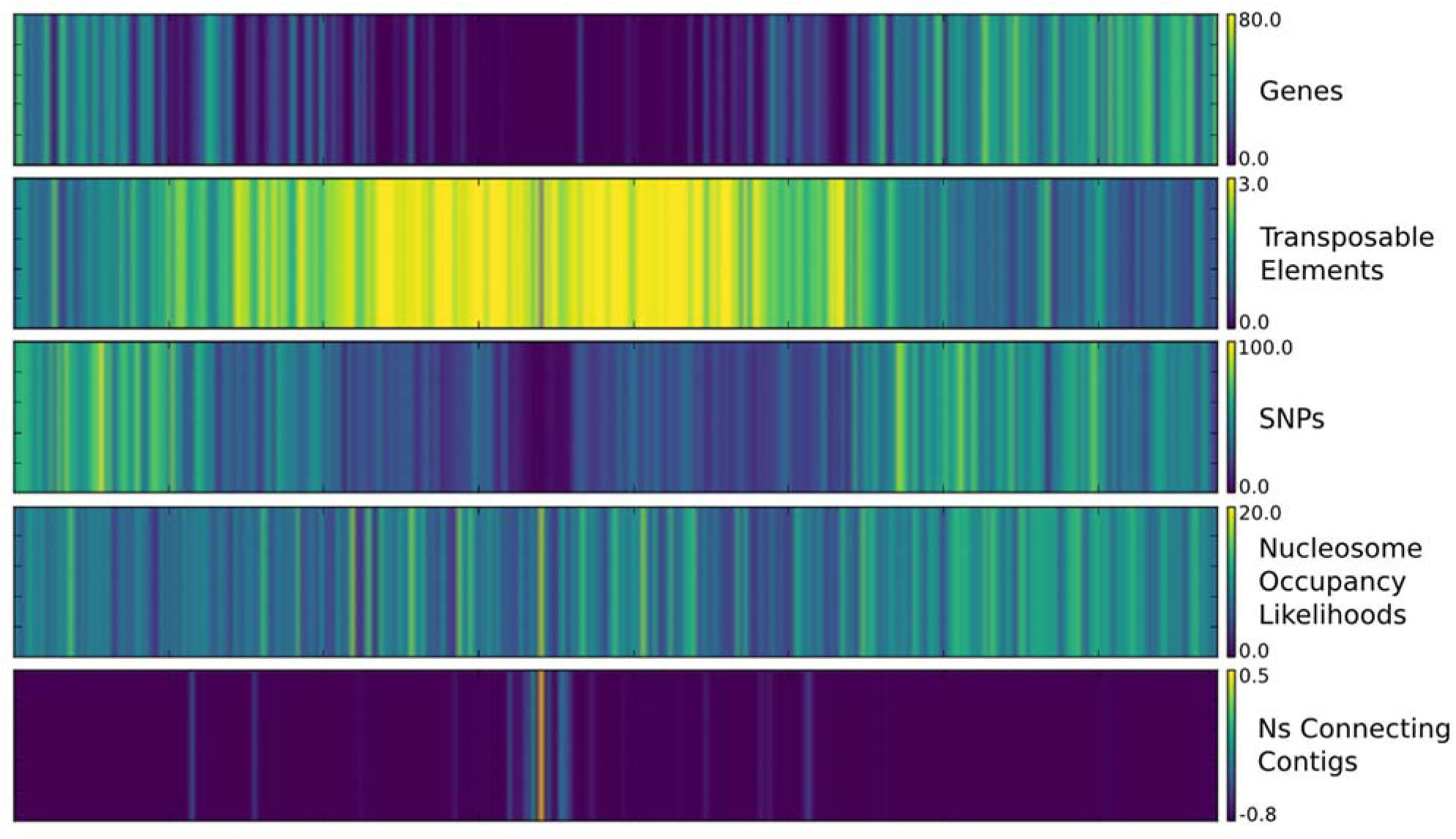
Feature densities and score averages across chromosome 2 of the sorghum genome. Color map displaying the average densities of multiple features across chromosome 2 of the sorghum genome, including annotated genes, transposable elements, single nucleotide polymorphisms, nucleosome occupancy likelihoods, and uncalled bases (Ns) connecting contigs in the assembly. Maps for all 10 chromosomes are depicted in Supplemental Figures S1 and S2.

The distributions of genes, repetitive elements, and genetic variants across each sorghum chromosome were generated using 1Mbp sliding windows (Figure 2, Supplemental Figures S1 and S2). Genes are at higher density in the distal euchromatic regions of chromosome arms and repetitive sequences related to transposable elements are most dense in heterochromatic pericentromeric regions characteristic of sorghum chromosomes (Evans et al., 2013; Paterson et al., 2009). The accumulation of genetic variation in *Sorghum bicolor* accessions was examined by aligning and comparing reads from 56 resequenced sorghum genotypes to the v3 genome sequence. *Sorghum propinquum* samples and two subsp. *verticilliflorum* genotypes were removed before analyses of variant distribution due to their evolutionary divergence from BTx623 and other resequenced *Sorghum bicolor* genotypes. The analysis identified 7,375,006 single nucleotide polymorphisms (SNPs) and 1,876,974 insertion/deletions (indels) distributed across the 10 chromosomes. The density of genetic variants was highly variable across the sorghum genome, with higher variant density in the distal euchromatic regions relative to heterochromatic pericentromeric regions of each chromosome, consistent with previous reports (Evans et al., 2013).

Predicted nucleosome positioning in the BTx623 v3 reference genome was examined by generating nucleosome occupancy likelihoods using a support vector machine trained on human chromatin data and validated in maize. Using this approach every nucleotide position was assigned a nucleosome occupancy likelihood (NOL) based on the primary sequence identity of a 50 bp window centered on the nucleotide (Fincher et al., 2013; Gupta et al., 2008). While primary sequence is not the only determinant of nucleosome binding, it influences the relative affinity of binding and general trends are indicative of chromatin organization. The predicted nucleosome occupancy likelihoods for sorghum are similar to maize in that the distributions vary across each chromosome, but with a relatively uniform pattern that does not match variation in gene or repeat density across each chromosome (Figure 2, Supplemental Figures S1 and S2).

### 4.3 Periodicity in features related to variant distributions in the sorghum genome

Information in eukaryotic genomes is stored at multiple scales, ranging from single base-pairs that specify codon identity to megabase-sized topologically associated domains that regulate transcriptional states (Bonev and Cavalli, 2016). Some of these organizational properties are correlated with periodic signatures in the accumulation of genetic variation. For example nucleosome positioning generates periodicity in the accumulation of genetic variants in humans and medaka (Higasa and Hayashi, 2006; Sasaki et al., 2009; Tolstorukov et al., 2011). Given that these organizational properties are associated with genomic signals such as variant density, digital signal processing techniques can be used to identify signatures associated with these properties. To this end, the Discrete Fourier Transform (DTF) was used to examine periodicities in the accumulation of genetic variation and nucleosome occupancy likelihoods to help identify mechanisms by which the sorghum genome stores information.

A known functional feature of the genome that influences the accumulation of genetic variation is the wobble base in codons. Due to redundancy in the genetic code, every third base downstream of a coding sequence start site is under relaxed selection since the primary DNA sequence is often able to change without dramatically influencing the information content of the sequence. This manifests as a prominent periodicity with a period of 3 bp after processing the polymorphism accumulation signal in the coding sequence of sorghum genes for regions downstream of coding start sites, but not upstream (Figure 3).

**Figure 3:**
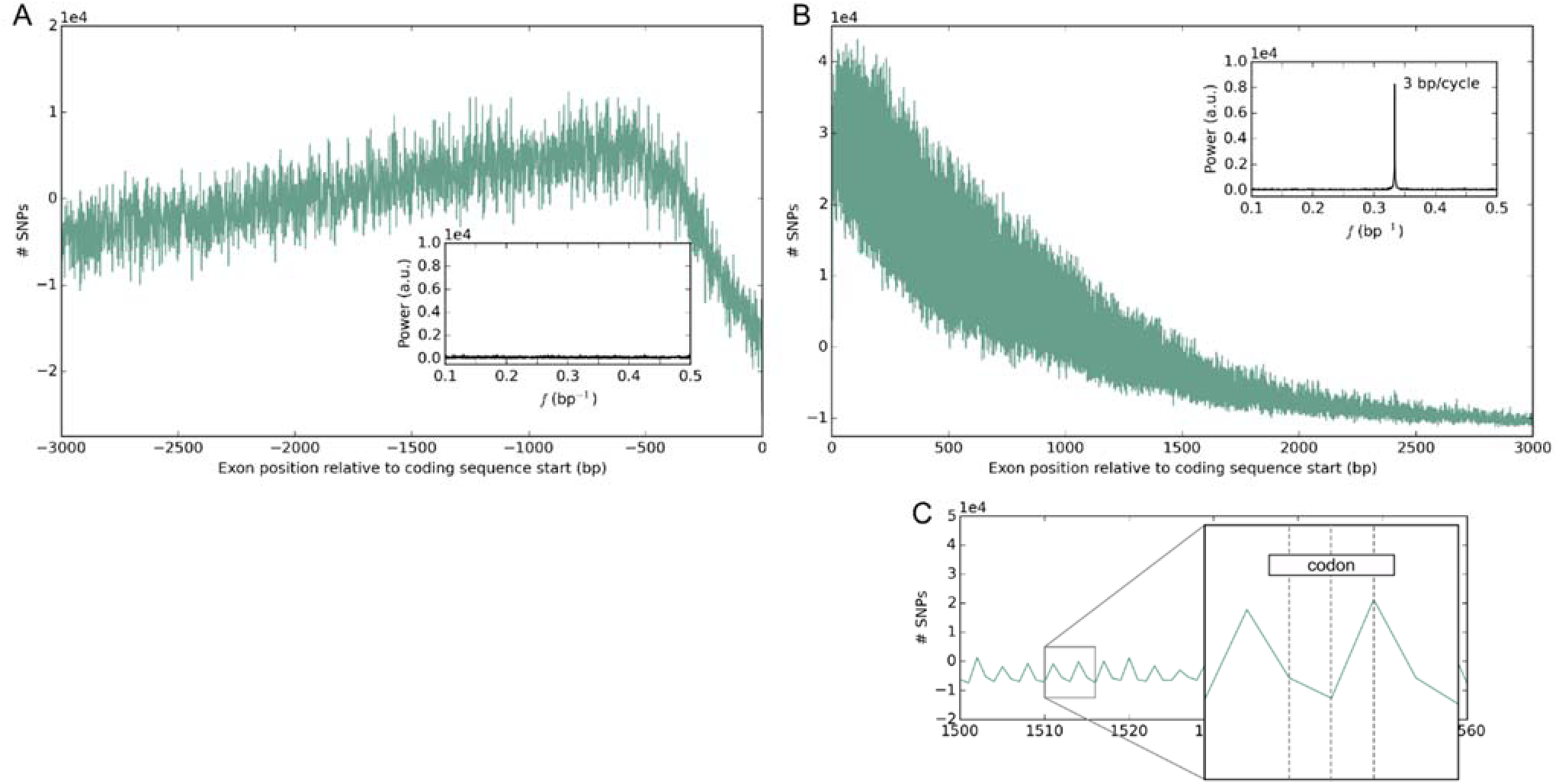
Functional properties of the sorghum genome leave periodic signatures that can be identified using signal processing techniques. Due to the degeneracy of the genetic code, relaxed selection at the wobble base in codons causes SNPs to accumulate with a periodicity of 3 bp downstream of coding sequence start sites in exon sequence (B), but not upstream of coding sequence start sites in the sorghum genome (A). This manifests as a strong signal at 0.33 bp-1 after transforming the SNP accumulation signal with the DFT (inset of B). (C) Zoom in of panel B shows the periodic signal. The Y axis of panels A and B plot the number of SNPs relative to the average of the respective window. The Y axis represents the sum of SNPs at each position relative to the CDS start site across all genes in the genome, centered to the mean of the respective window; CDS lengths of less than 3000 were considered to have 0 SNPs between their end and 3000 bp, leading to the apparent decline observed in panel B.

Nucleosome scale variant periodicities were examined for signatures of genome organization because studies in medaka and human indicated that genetic variation accumulates at transcription start sites (TSSs) with periodicities around 150 bp, corresponding to nucleosome occupancy (Higasa and Hayashi, 2006; Sasaki et al., 2009). To determine if a similar phenomenon was present in the sorghum genome, the genetic variation that accumulated around transcription start sites as well as nucleosome occupancy likelihoods were examined. Consistent with micrococcal nuclease digestion results in maize and *Arabidopsis,* prediction scores indicated a high likelihood of a nucleosome positioned immediately downstream of the transcription start site of genes in sorghum (Figure 4) (Fincher et al., 2013; Liu et al., 2015). While variant frequency decreased immediately downstream of TSSs, the variant profile in sorghum did not show accumulation of genetic variants with a period of ~150 bp downstream of these sites. Nucleosome occupancy predictions also did not predict a periodic arrangement of nucleosomes downstream of transcription start sites.

**Figure 4:**
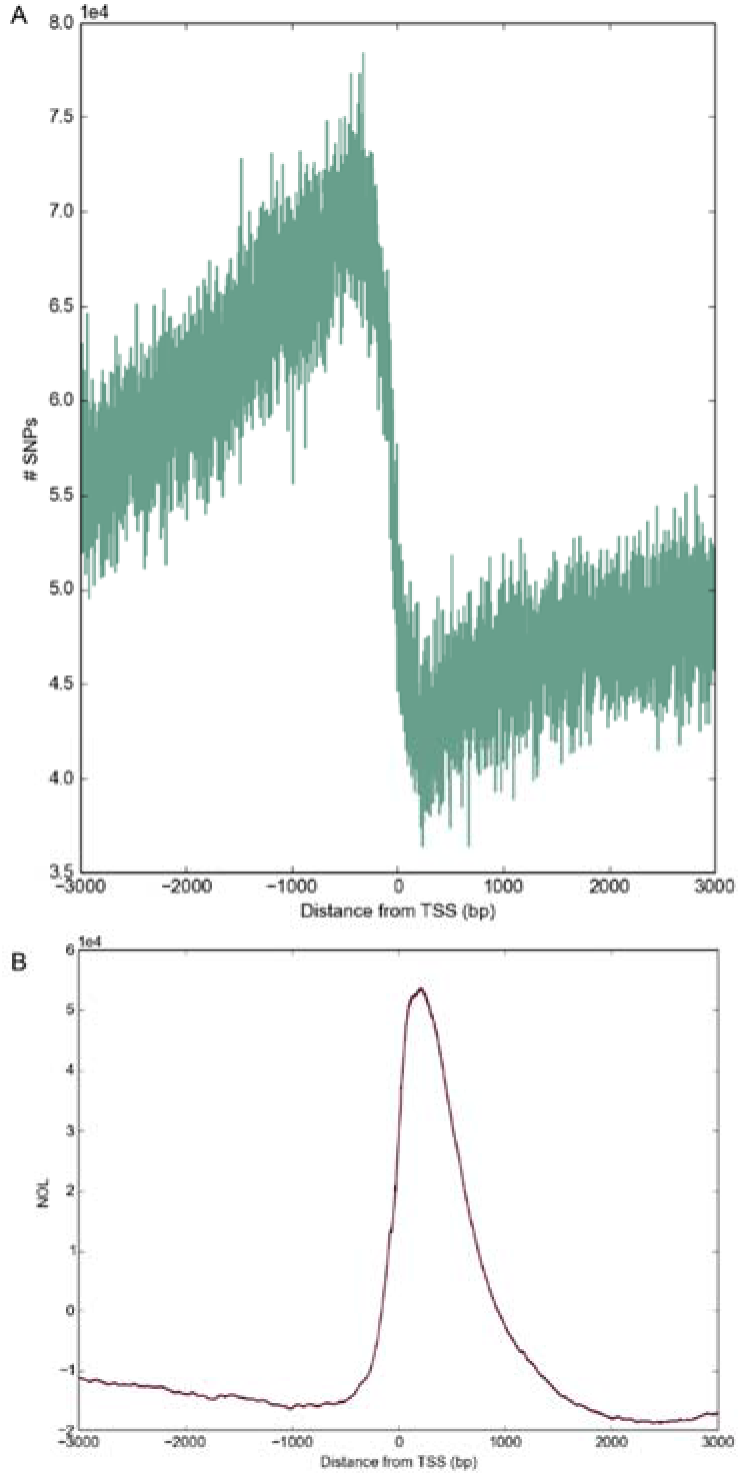
Genetic variation and nucleosome occupancy likelihoods around transcription start sites in the sorghum genome. Nucleosome occupancy scores indicate a high likelihood of a nucleosome positioned immediately downstream of transcription start sites in sorghum. Strong evidence that nucleosomes were stably positioned based and periodically arrayed as in medaka and human was not observed in either the accumulation of genetic variants nor nucleosome occupancy likelihoods, though NOLs indicate that a nucleosome is often positioned immediately downstream of the transcription start site, consistent with experimental observations in maize and *Arabidopsis*.

Nucleosome scale periods of 180 bp are present in nucleosome occupancy likelihood profiles in multiple regions of the genome, and are especially pronounced in subtelomeric regions, suggesting the possibility of stably positioned, periodically arrayed nucleosomes downstream of the (CCCTAAA)_n_ telomere repeats present at the end of sorghum chromosomes (Figure 5B and 5C) (Klein et al., 2000).

**Figure 5:**
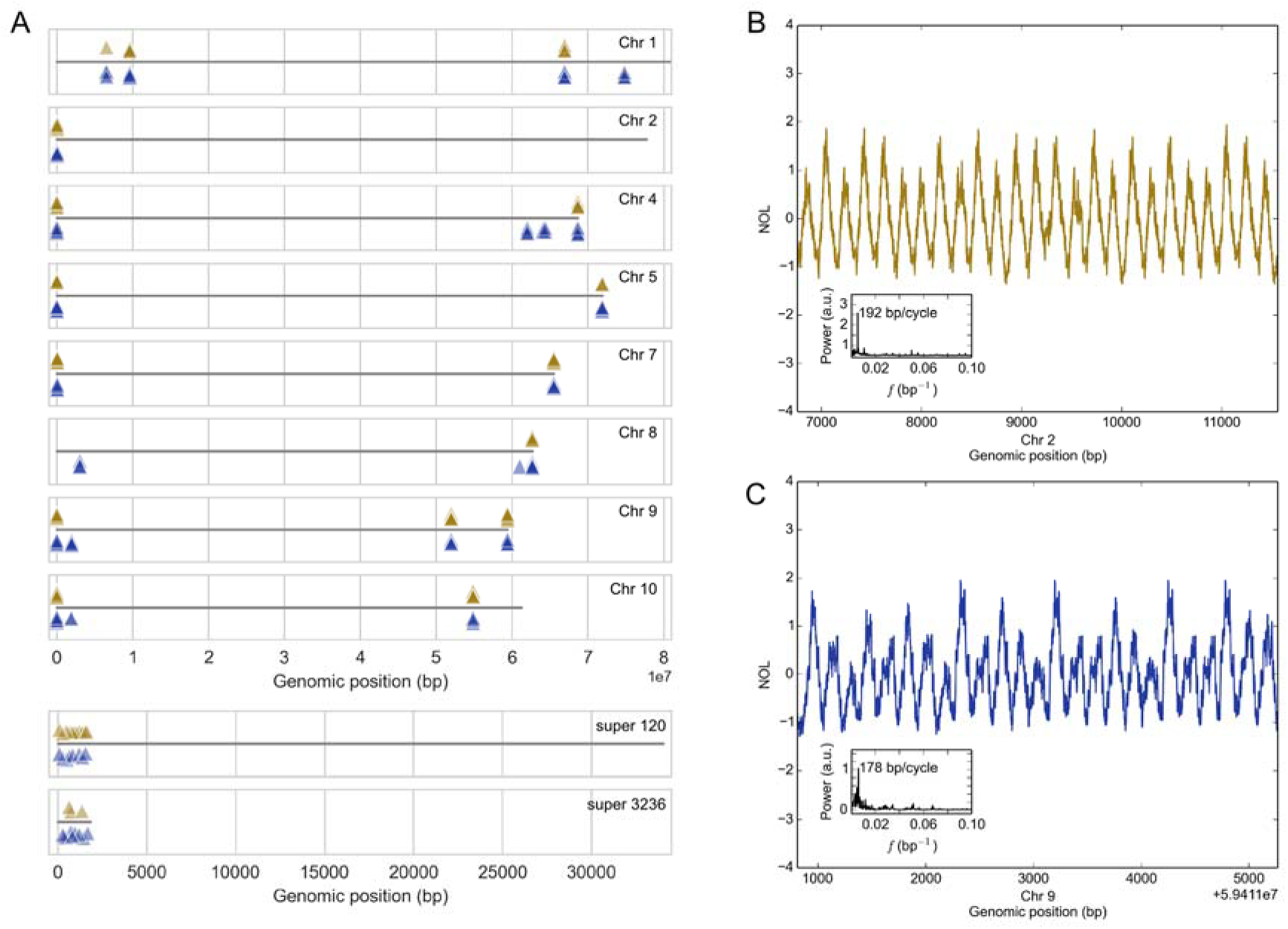
Subtelomeric periodicities in nucleosome occupancy likelihoods correspond to arrays of tandem repeats located near the end of most chromosome arms. (A) Graphic representation of BLAST hits for the consensus sequence of STA1 and STA2 indicate that most chromosome arms contain subtelomeric tandem arrays of the STA1 or STA2 monomer; two super contigs in the assembly also contain arrays, and may correspond to subtelomeric sequence on the arm of chromosome 2. (B) Nucleosome occupancy likelihoods (centered on the mean) and power spectrum for an array of the STA1 monomer with multiple sequence alignment of continuous arrays from multiple chromosome arms. (C) Same as panel A, but with arrays of the STA2 monomer. STA1 and STA2 share sequence identity and are likely related, though most chromosome arms bear tandem arrays of only one or the other; BLAST hits show colocalization due to shared identity.

Since the SVM used for nucleosome occupancy likelihood calculation used only primary sequence, any primary sequence that was tandemly arrayed (e.g., satellite DNA) should also yield a periodic signal. Further characterization of the primary sequence underlying the periodic signal identified that the periodicity indeed resulted from tandemly arrayed, subtelomeric, satellite DNA with a repeat size of 180 bp, consistent with observations that the monomer size of satellite DNA repeats often correspond to the length of DNA wrapped around nucleosomes (Mehrotra and Goyal, 2014). BLAST analyses indicated that most chromosome arms contained tandem arrays of one of two satellite repeats, with the two types of repeats sharing some sequence identity (Figure 5A). The two monomers are referred to as subtelomeric tandemly arrayed 1 and 2 (STA1 and STA2) here for brevity.

Tandem arrays of STA1 or STA2 (or a complex mixture of both) exist on most of the sorghum chromosome arms, with the longest array present at the beginning of chromosome 2, repeating STA1 more than 200 times over more than 36 kbp. Arrays of STA1 or STA2 are present within 50 kbp of the beginning and end of chromosomes 4, 5, 7, and 9. Chromosomes 3 and 6 are the only scaffolds without the elements near the ends of one of the chromosome arms (Figure 5). Notably, the arrays are also found on super contigs 120 and 3236; these may correspond to the ends of one or more chromosomes, although they lack the (CCCTAAA)_n_ telomeric repeat. Telomeric repeats were found at both termini of chromosomes 1, 4, 5, 7 and 10 and at one of the two termini of chromosomes 2, 3, 6, 8 and 9, so no strong relationship between the presence of an assembled telomere and the STA repeat was observed (Supplemental Table S1).

Alignment searches for STA1 and STA2 in maize, rice and more distantly related plants suggest that this sequence repeat feature is sorghum specific. *De novo* repetitive element annotation identified the arrays as individual terminal-repeat retrotransposons in miniature (TRIM) elements, although they were not included in a recent annotation of plant TRIMs, a database that includes sorghum (Gao et al., 2016). While TRIMs have been observed to accumulate in tandem arrays, the monomers of STA1 and STA2 lack most of the features of canonical TRIM elements (Gao et al., 2016; Witte et al., 2001). Only STA1 bears a putative primer binding site (PBS; complementary to the sorghum methionine tRNA). Notably, STA1 shares sequence identity with an unclassified sorghum element (SRSiOTOT00000007) from the TIGR Plant Repeat Database (Ouyang and Buell, 2004), as well as the *S. halepense*-specific repetitive elements XSR6, XSR1, and XSR3 (Hoang-Tang et al., 1991). The STA1 and STA2 monomers both have a complex substructure of internal duplication and tandem repeats (Figure 6, Supplemental Figure S3, and Supplemental File S2).

**Figure 6:**
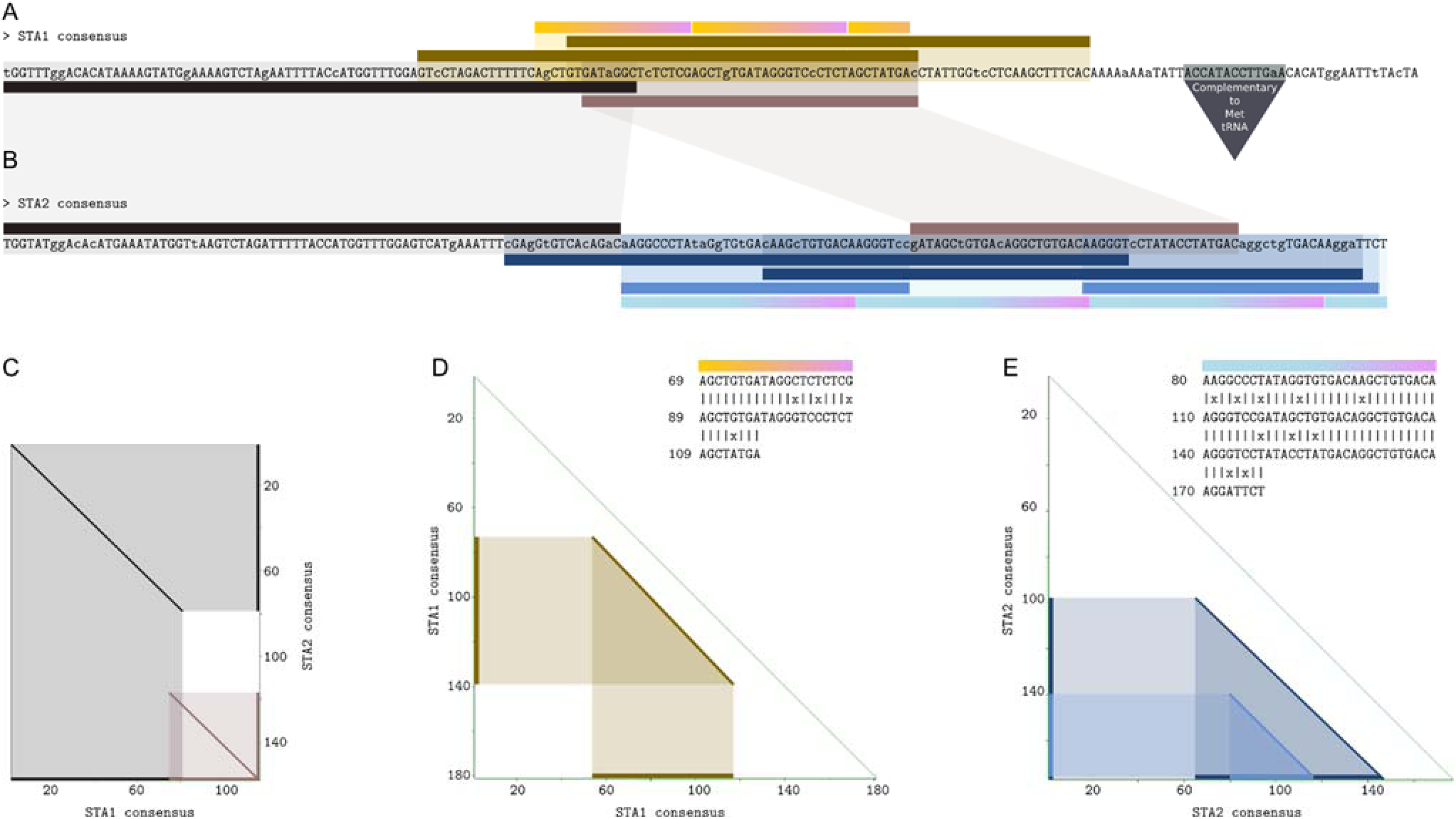
Relationships and structure of STA monomers. Monomers of STA1 and STA2 share blocks of identity and each contain direct repeats. (A, B) Consensus sequence of the STA1 and STA2 monomers annotated for sequence shared between monomers as well as direct repeats within each monomer. Annotations are indicated by colored bars and color coded according to their origin, either from (C) an alignment dotplot of STA1 and STA2, (D) a self-alignment dotplot and tandem direct repeat analysis of STA1, or (E) a self-alignment dotplot and tandem direct repeat analysis of STA2. STA1 also contains a putative promoter binding site (PBS) complementary to a sorghum methionine tRNA, a feature characteristic of TRIM elements (Witte et al., 2001).

Signatures generated as a consequence of genome regulation or structure can be detected by signal processing techniques as shown previously for the effects of the wobble position in codons (Figure 3). Therefore, variant accumulation at larger scales was examined across the genome to identify periodic patterns of SNP accumulation. Consistent with previous reports, DNA variant profiles obtained from comparison of sorghum genome sequences from genotypes representing different sorghum races revealed that variants accumulate in a non-uniform fashion across the genome (Evans et al., 2013). Genome-wide analysis of SNP accumulation using the DFT indicated that this non-uniformity occasionally manifested as a periodic event where genetic variants accumulated in peaks and troughs in a region of the genome. Genome-wide scans of variant accumulation using the DFT indicated that multiple regions of the genome display large-scale periods, such that a peak of variant accumulation is observed every 25 kbp (Figure 7). As with the periodicity observed at the wobble base, the cyclical nature of peaks in variant accumulation may represent a consequence of genome organization or information storage. This large scale periodicity of SNP accumulation was observed in regions of chromosomes 1, 3, 4, 5, 9, and 10 when SNPs called from sequence data for 52 sorghum genotypes were analyzed.

**Figure 7:**
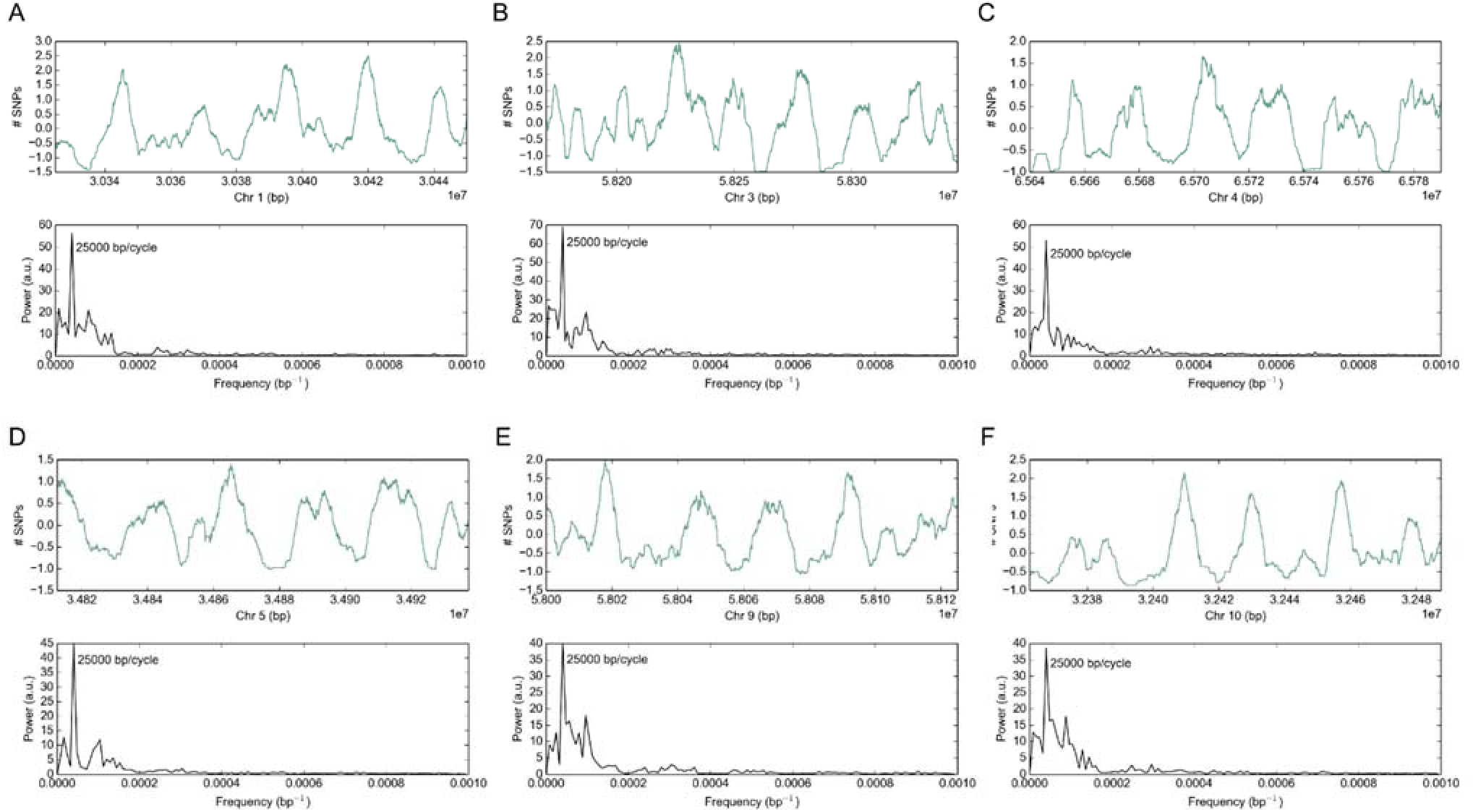
Periodicities in the accumulation of genetic variation in the sorghum genome. A genome-wide scan for periodic accumulation of SNPs identified multiple regions of the genome with a distinct period of 25,000 bp. The top plot of each panel shows the accumulation of SNPs relative to the mean of the window given a 5,000 bp sliding average, and the bottom plot shows the power spectrum after transformation with the Discrete Fourier Transform (A-F).

### 4.4 The Sorghum Transcriptome Atlas

The sorghum transcriptome atlas used to improve the sorghum reference genome gene annotation represents a broad diversity of tissues, developmental stages, and responses to nitrogen sources, encompassing a variety of transcriptional states. The transcriptome atlas was developed with two primary goals: (1) to sample the major plant organs (roots, leaves, stems, panicles) when these organs were growing and then following maturation at different developmental stages (juvenile, vegetative, reproductive) to facilitate comprehensive annotation of genes in the sorghum genome and (2) to sample a diversity of nitrogen states and sources as part of an inter-species plant gene atlas project. A thorough analysis of these datasets is beyond the scope of this manuscript, but they are described here for release into the public domain for use by the community at large. The samples collected are described in Supplemental Table S2 and Supplemental File S3.

Initial analyses of the transcriptome data were carried out to provide a high-level overview of the transcriptome atlas contents. Correlations of the expression values across all 34,211 genes indicated high correlation within biological replicates of the same sample, as well as correlated groups between samples from the same tissue (Supplemental Figure S4). The largest block of correlated expression was a block of high correlation between all of the root samples, regardless of whether the root sample was more distal or proximal or root nitrogen treatment. Dormant seed shared the least correlation with any of the samples, indicating that its steady state pool of transcripts differed the most dramatically from other tissues analyzed.

Hierarchical clustering based on the transcript abundance of all 34,211 genes via UPGMA identified similar relationships among the samples, indicating that the transcript pool of a given sample was defined predominantly by the tissue/organ identity rather than the developmental stage. Seed samples were the most transcriptionally distinct, especially dormant seed. In agreement with hierarchical clustering, k-means clustering indicated that roots, stems, leaves, and seeds formed distinct clusters based on gene expression (Figure 8).

**Figure 8:**
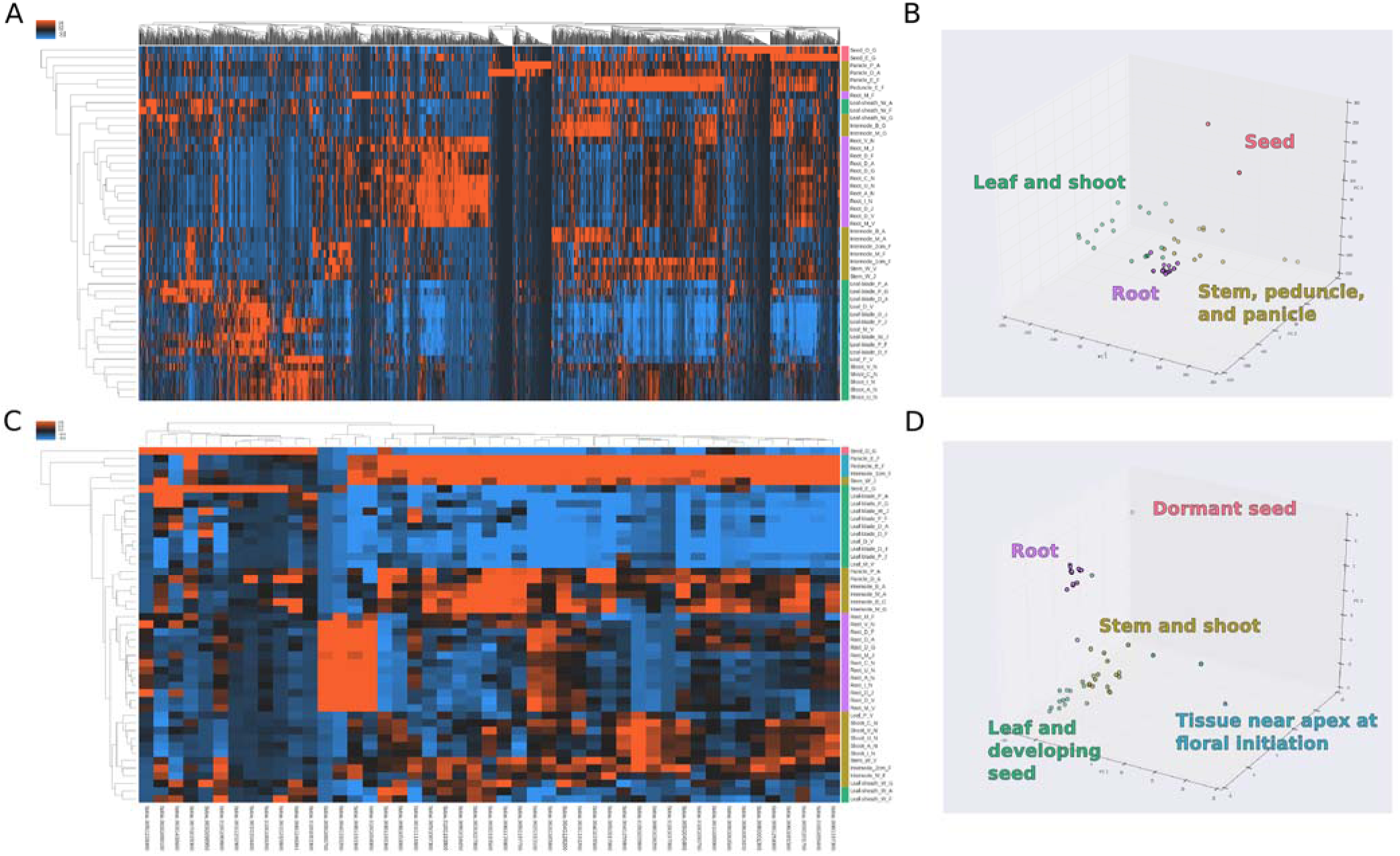
Clustering and ontological analyses indicate the expression of kinase genes are associated with tissue identity. (A) Heat map and hierarchical clustering of atlas samples based on gene expression of all 34,211 sorghum genes; color bars on right correspond to k-means clusters in panel B. (B) Scores of the first three principal components of the atlas samples colored based on k-means cluster (k = 4) using expression values of all genes. (C) Ontological enrichment analysis of the 2,500 genes with the largest loadings for the first three principal components indicate that kinase genes were overrepresented, and the expression of the 47 kinase genes driving the enrichment are plotted as a heat map with hierarchical clustering; color bars on the right correspond to k-means clusters in panel D. (D) Scores for the first three principal components of the atlas samples colored based on k-means cluster (k = 5) using expression values of the 47 kinase genes.

To identify a set of genes with large variation in expression across the dataset, principal component analysis was performed using all 34,211 genes to obtain the first three principal components (PCs), and the set of 2,500 genes with the largest sum magnitude of loadings for the first three PCs were identified (Supplemental Figure S5). To determine if particular classes of genes were overrepresented among these genes that explained large components of variation in the dataset, ontological enrichment analysis was performed for terms related to molecular functions. Three molecular function enrichments were identified, including structural constituents of the ribosome (GO0003735), protein kinase activity (GO0004672), DNA-directed RNA polymerase activity (GO0003899) (Supplemental File S4). Since kinase activity is associated with signal transduction and the other two terms may be explained by the developmental state of the tissue (e.g., high transcription and translation activity), the 47 genes responsible for protein kinase activity were investigated further (Supplemental Table S3).

Hierarchical clustering and k-means clustering based on the expression state of the 47 kinase activity genes broadly reproduced the same groupings as all 34,211 genes (Figure 8). Notably, three samples associated with proximity to the shoot apical meristem at floral initiation were clustered together, potentially representing kinases involved in the transition from a shoot apical meristem to a floral meristem.

## 5 DISCUSSION

The sorghum genome sequencing project was organized in 2005 because *Sorghum bicolor* has a relatively simple genome compared to many other grasses, sorghum is a valuable genetic model for C4 grass research, and sorghum crops are important world wide, especially as subsistence crops in the semi-arid tropics (Participants, 2005). The sorghum genome sequence improved our understanding of sorghum genome organization, coding capacity, and aided analysis of grass genome diversification (Paterson et al., 2009). The original reference genome sequence was based on ~8.5- fold depth paired-end Sanger sequence reads from genomic libraries with 100-fold variation in the size of inserts. Discrepancies in order that arose during assembly were resolved in part using information from pre-existing high resolution genetic and BAC-based physical maps of the sorghum genome (Bowers et al., 2003; Klein et al., 2000). The sum of the 201 largest sequenced scaffolds spanned 678.9 Mbp of which 625.7 Mbp of the sequence was assigned chromosomal locations. The size of the reference genome had been previously estimated by flow cytometry to be 818 Mbp (Price et al., 2005), indicating that reference genome v1 sequence comprising the 10 chromosomes accounted for ~76% of the total genome sequence. It was reported that 15 of the 20 chromosome termini contained telomeric repeats and that Cen38 sequences (Zwick et al., 2000) were present in each chromosome, although these sequences were also found in many of the sequence scaffolds that could not be incorporated into the chromosomal sequences (Paterson et al., 2009). Despite the need for further improvement, the resulting sorghum reference sequence has been of great value to the sorghum and grass research community, enabling comparative genomics (Paterson et al., 2009), association studies (e.g.,Brenton et al., 2016; Morris et al., 2013), the development of genotyping by sequencing methods for sorghum (Morishige et al., 2013), analysis of sorghum diversity and variant distribution (Evans et al., 2013; Mace et al., 2013; McCormick et al., 2015), genome methylation profiles (Olson et al., 2014), and many other research activities.

The objective of the current study was to update the sorghum reference genome sequence and its annotation, and to characterize additional features of the sorghum genome that affect sorghum biology. The sequence quality and coverage of the reference genome was improved by obtaining 110X coverage of the genome using Illumina sequencing, targeted finishing of ~344 Mbp of gene rich portions of the genome, and by improving order and sequence contiguity using a high density genetic map. These activities increased sequence coverage by ~30 Mbp, reduced error frequency 10- fold to ~1/100 kbp, and improved assembly order by moving a 1 Mbp block of DNA from SBI-06 to SBI-07. The research did not identify and incorporate sequences containing telomeric repeats that are missing from the ends of 5 chromosomes and the order and completeness of sequences in the pericentromeric regions that have high repeat density was not significantly changed. Long read sequencing and Hi-C analysis (Sanborn et al., 2015) would be valuable approaches to implement to further improve the reference genome sequence.

Version 1.4 of the sorghum genome sequence provided evidence for 27,604 annotated genes. Subsequent analysis of gene annotations that incorporated RNA-seq data indicated that a large number of genes were not annotated in v1.4 and that many of the annotations were incomplete (Olson et al., 2014). Results from the current study based on deep RNA-seq analysis of 47 tissues from roots, stems, leaves, leaf sheaths, panicles and seed enabled the annotation of 34,211 genes, a 24% increase relative to v1.4. RNA-seq data also improved the annotation of exons resulting in a significant increase in average gene size consistent with prior results based on a similar approach (Olson et al., 2014). Increased gene coverage and improved gene annotation and sequence accuracy will aid comparative genomics studies as well as GWAS and map-based QTL to gene discovery projects that can result in false negatives/positives if the reference genome sequence used for analysis is not a well annotated high quality sequence. In our own research, errors and misannotation of the v1 sequence caused identification candidate gene alleles underlying QTL to be missed until direct sequencing was carried out on all genes in fine mapped intervals (Hilley et al., 2016; Murphy et al., 2011).

While v3.1 is a substantial improvement over v1, additional information is needed to fill in missing portions of the genome sequence and to improve gene annotation. As noted above, one end of 5 chromosomes lack telomeric sequences indicating these chromosome sequences are not complete. Moreover, it is likely that the sequence of the pericentromeric repeat-rich regions of chromosomes is incomplete and possibly misordered in some regions. Since recombination is extremely low across the large heterochromatic pericentromeric regions (Kim et al., 2005), the high resolution genetic map employed to order DNA in euchromatic regions was not useful for ordering sequences across the pericentromeric regions. A combination of long range, long read sequencing and Hi-C analysis would be useful to improve these regions of the reference genome. In addition, Iso-Seq was shown to aid the analysis of full-length splice isoforms, alternative polyadenylation sites, and non-coding RNAs in sorghum (Abdel-Ghany et al., 2016). The analysis showed that in depth Iso-Seq data will significantly improve the current annotation of the sorghum genome and transcriptome. Moreover, pan-genome projects in maize and other species show that a substantial number of ‘dispensible’ genes are found only in a subset of the genotypes of a species germplasm (Hirsch et al., 2014). Therefore characterization of the sorghum pan-genome will require the acquisition and de novo assembly of genomes from diverse sorghum genotypes possibly aided by the construction of a set of reference genomes sequences that sample sorghum’s diversity space.

The distribution of genes, repeats, variants, and other features of the sorghum genome was updated based on the v3 genome sequence. Gene density was highest in distal euchromatin portions of chromosomes and repetitive sequences related to retrotransposons were enriched in heterochromatic pericentromeric regions as previously described (Kim et al., 2005; Paterson et al., 2009). Predicted nucleosome positioning based on primary sequence data showed localized variation in nucleosome density but a fairly uniform distribution of nucleosome localization across chromosomes. Digital signal processing of genomic signals is a useful approach to identify novel patterns in genome structure. Through this approach, previously uncharacterized subtelomeric tandem repeats were identified in sorghum. The importance of satellite DNA in influencing plant genome organization has been documented previously, and subtelomeric tandem arrays are characteristic of many plant genomes, raising the possibility that they play a role in telomere or genome stability (Mehrotra and Goyal, 2014; Padeken et al., 2015).The subtelomeric repeats STA1 and STA2 were located near the distal ends of most chromosomes. These sequences were identified as TRIM-like, although they lacked most of the sequence motifs found in TRIMs identified in other plants (Gao et al., 2016; Witte et al., 2001). The function of these subtelomeric repeats is unknown, although subtelomeric repeats have been shown to be involved in bouquet formation and to facilitate the pairing of homologous chromosomes during meiosis (Harper et al., 2004; Sadaie et al., 2003). A complete analysis of these subtelomeric arrays will require additional long-read sequencing to fully characterize the size and location of these subtelomeric repeats and to determine if they are present in all of the sorghum chromosomes.

Comparison of whole genome sequences from 52 diverse sorghum genotypes to the v3 reference genome sequence identified ~7.8M SNPs and ~1.9M indels. Large scale signals in the accumulation of genetic variation were identified by signal processing techniques, and these may represent signatures left by higher order organization. For example, elevated variant frequency was associated with the wobble position in codons. Previous studies had documented elevated variant density in euchromatic regions compared to pericentromeric regions of sorghum chromosomes and significant variation in variant density within euchromatin when the genomes of different sorghum races were compared (Evans et al., 2013). Genetic hitchhiking may be acting to reduce genetic variation in regions of low recombination near centromeres (Barton, 2000). In this study, variant distributions based on the analysis of 52 sorghum genomes were analyzed and found to contain large scale variant distribution features that repeat every ~25 kbp. We had previously speculated that large scale features like these could be generated by regional variation in recombination and repair, possibly due to higher-order chromatin organization (Evans et al., 2013). In addition, the ability of DNA repair machinery to access and correct mutations and selection pressures generated by functional properties of the genome such as gene coding sequences across the gene rich distal arms of sorghum chromosomes where rates of recombination are high could be influencing the accumulation of variants (Evans et al., 2013; Mace et al., 2013; Makova and Hardison, 2015; Zheng et al., 2011). Additional analyses should leverage wavelet transforms in addition to the discrete Fourier transform to resolve problems associated non-stationary signals, as these genomic signals are likely non-stationary in nature. Moreover, wavelet transform coefficients can be used to correlate multiple features such as recombination and genetic variation (Spencer et al., 2006). The results from digital signal processing approaches used to examine the sorghum genome indicate that additional experimentation to annotate sorghum chromatin as well as higher order features like chromatin interactions and nuclear lamina binding sites will be useful to better understand factors shaping the landscape of the sorghum genome.

The RNA-seq transcriptome atlas reported here focused on the collection of tissue from growing and fully developed roots, stems, leaves, panicles and seeds during development. Collection started with seed germination, traversed the juvenile, vegetative and reproductive phases concluding with the analysis of the transcriptome of dry seed. This transcriptome atlas complements prior RNA-seq data collected from sorghum stems during 100 days of development that included the phase of sucrose accumulation (McKinley et al., 2016), sorghum transcriptome responses to dehydration and ABA (Dugas et al., 2011), dynamic changes in tiller bud transcriptomes modulated by PhyB (Kebrom and Mullet, 2016), and an analysis of meristematic tissues, florets, and embryos (Olson et al., 2014). An in depth description of the RNA-seq data is underway, however results described here show that the atlas is of high quality and useful for the analysis of tissue and developmental states. The expression of genes encoding kinases was found to differentiate transcriptome tissue states identified by PCA analysis. Kinases are involved in plant development and tissue identity, and the transcriptome atlas identified 47 genes encoding kinases whose transcript abundance broadly distinguishes between tissue types. The kinase genes represent putative regulators of tissue identity in sorghum, and some were previously characterized to influence plant development. Among the intersection of kinases identified from the sorghum transcriptome atlas and those previously characterized in the literature include kinases like WAK2, which is required for cell expansion during development by monitoring pectin (Kohorn, 2015). TSL mediates RNAi silencing and may influence development (Uddin et al., 2014). WNK4 and WNK6 were found to be regulated by the circadian clock and may be involved in regulating flowering time (Nakamichi et al., 2002; Wang et al., 2008). ACR4 is associated with maintenance of root stem cell identity in the RAM with CLV4, though ACR4 was not expressed in roots in the transcriptome atlas (Stahl et al., 2013). ERL2 controls organ growth and flower development via cell proliferation (Bemis et al., 2013; Shpak et al., 2004). YODA influences root development through auxin up-regulation and cell division plane orientation (Smékalová et al., 2014). These represent a small sampling of putative regulators of sorghum development, and thus the sorghum transcriptome atlas represents a valuable resource with which to both annotate the sorghum genome and to promote characterization of the gene regulatory networks underlying sorghum development.

## 6 METHODS

### 6.1 Genome assembly and improvement

320 regions of the version 1 sorghum reference genome assembly (Paterson et al., 2009) that contained a gene density greater than 2 genes per 100 kb were chosen for finishing. Finishing was performed by resequencing plasmid subclones and by walking on plasmid subclones or fosmids using custom primers. Small repeats in the sequence were resolved by transposon-hopping 8 kb plasmid clones, while 454 and Illumina based small insert libraries were used to improve resolution of simple sequence repeats. To fill large gaps, resolve large repeats, or to resolve chromosome duplications and extend into chromosome telomere regions, complete fosmid and BAC clones were shotgun sequenced and finished. The finished sequence was assembled, and each assembly was validated by an independent quality assessment. Finished regions were integrated by aligning the regions to the existing V1.0 assembly. 349 regions representing 344.4 Mbp of sequence were integrated in this manner.

A high-density genetic map generated from 437 recombinant inbred lines from a cross of BTx623 and IS3620C was used to improve the quality of the assembly and increase its coverage by integrating additional sequence scaffolds (Burow et al., 2011; Truong et al., 2014) into the 10 linkage groups. Scaffolds were broken if they contained a putative false join coincident with an area of low BAC/fosmid coverage. A total of 8 breaks were identified in the V1.0 release chromosomes, and an additional 7 previously unmapped scaffolds were integrated into the assembly in the appropriate location (Supplemental File S1). A 1.08 Mb region of the V1.0 chromosome 6 was moved to chromosome 7. 15 joins were made to form the final assembly containing 10 chromosomes capturing 655.2 Mb (97.1%) of the assembled sequence. Each join was padded with 10,000 Ns.

Homozygous variants identified from 110x of 2x250 (800 bp insert) Illumina fragments sequenced from the same DNA isolation as the original sequence were obtained and used to correct sequencing errors in the reference assembly. Reads were aligned to the integrated assembly and variants were called; variants that were called as homozygous were considered as candidates for correction in the reference assembly. A total of 1,942 (41% of called) homozygous SNPs and 1,432 (82% of called) homozygous indels were corrected in the process. SNPs and/or INDELs that were within 150bp of one another were not corrected. Additional information regarding methods of assembly and finishing are contained in Supplemental File S1.

### 6.2 Sample preparation and sequencing for transcriptome atlas and whole genome resequencing.

The reference line BTx623 was grown under 14 hour day greenhouse conditions in topsoil, equivalent to native field soil from Brazos County, TX, to generate tissue for two separate experiments: (1) a tissue by developmental stage timecourse, and (2) a nitrogen source study. For the tissue by developmental stage timecourse, plants were harvested at the juvenile stage (8 DAE), the vegetative stage (24 DAE), at floral initiation (44 DAE), at anthesis (65 DAE), and at grain maturity (96 DAE) and leaf, root, stem and reproductive structures were flash frozen in liquid nitrogen. For each tissue by stage combination, three biological replicates (i.e. three plants representing a single condition) were harvested with the exception of the juvenile stage, for which a replicate was represented by five plants instead of one to compensate for lower tissue abundance. For the nitrogen source study, plants grown under differing nitrogen source regimes were harvested at 30 DAE, and shoots and roots were flash frozen. For each tissue by condition, three biological replicates were obtained. Additional details regarding harvested samples can be found in Supplemental Table S2 and Supplemental Files S1 and S3.

Tissue was ground under liquid nitrogen and RNA was extracted using a Trizol-reagent based extraction. Tissues with high levels of starch used a modified Trizol-reagent protocol (Li and Trick, 2005). Plate-based RNA sample prep was performed on the PerkinElmer Sciclone NGS robotic liquid handling system using Illumina’s TruSeq Stranded mRNA HT sample prep kit utilizing poly-A selection of mRNA following the protocol outlined by Illumina in their user guide: http://support.illumina.com/sequencing/sequencing_kits/truseq_stranded_mrna_ht_sample_prep_kit.html, and with the following conditions: total RNA starting material was 1 ug per sample and 8 cycles of PCR was used for library amplification. The prepared libraries were then quantified by qPCR using the Kapa SYBR Fast Illumina Library Quantification Kit (Kapa Biosystems) and run on a Roche LightCycler 480 real-time PCR instrument. The quantified libraries were then prepared for sequencing on the Illumina HiSeq sequencing platform utilizing a TruSeq paired-end cluster kit, v4, and Illumina’s cBot instrument to generate a clustered flowcell for sequencing. Sequencing of the flowcell was performed on the Illumina HiSeq2500 sequencer using HiSeq TruSeq SBS sequencing kits, v4, following a 2x150 indexed run recipe. Sequencing generated roughly 3.3 billion pairs of sorghum paired-end read data.

Nine additional sorghum lines (100M, 80M, BTx623, BTx642, Hegari, IS3620C, SC170-6-17, Standard Broomcorn, and Tx7000) were resequenced to supplement the 47 lines already available (Mace et al., 2013; Zheng et al., 2011). Seeds were soaked in 20% bleach for 20 minutes and washed extensively in distilled water for one hour. Seeds were germinated on water saturated germination paper in a growth chamber (14 hr light; 30° C/10 hr dark; 24° C). Genomic DNA was isolated from 8-day old root tissue using a FastPrep DNA Extraction kit and FastPrep24 Instrument (MP Biomedicals LLC, Solon, OH, USA), according to the manufacturer’s specifications. DNA template (350 bp average insert size) was prepared using a TruSeq^®^ DNA PCR-Free LT Kit, according to the manufacturer’s directions. Paired-end sequencing (125 x 125 bases) was performed on an Illumina HiSeq2500.

### 6.3 Transcriptome Annotation

The RNAseq reads were aligned to the updated reference assembly using GSNAP and assembled into 127,415 RNAseq transcripts with the PERTRAN pipeline (Shu et. al., unpublished). These transcripts were combined with 209,835 ESTs to generate 111,994 transcript assemblies using PASA. Loci were determined by transcript assembly alignments and/or EXONERATE alignments of proteins from *Arabidopsis thaliana*, rice, maize or grape genomes. Gene models were predicted by homology-based predictors, mainly FGENESH+, FGENESH_EST, and GenomeScan. The best scored predictions for each locus were selected using multiple positive factors including EST and protein support, and one negative factor: overlap with repeats. The selected gene predictions were improved by PASA by adding UTRs, splicing correction, and adding alternative transcripts. Finally, a homology analysis was performed on the PASA-improved models relative to the proteomes of *Arabidopsis thaliana*, rice, maize and grape to identify high quality gene models and remove models with extensive transposable element domains.

### 6.4 Additional feature annotation, feature coverage, and periodicity analyses.

Additional features were annotated in the sorghum genome, including repetitive sequence, genetic variants, and nucleosome occupancy likelihoods. Repetitive sequence, including transposons and SSRs, were annotated using both a de novo annotation and an annotation with existing libraries with REPET v2.5; existing repetitive element libraries included the TIGR Plant Repeat Database and RepBase (Bao et al., 2015; Flutre et al., 2011; Ouyang and Buell, 2004; Quesneville et al., 2005). Genetic variants were called from sequence data for 56 sorghum resquenced sorghum samples (Supplemental File S5). Processing of sequence reads to variant calls, including alignment to the Sbi3 reference genome, base recalibration, indel realignment, joint genotyping, and variant quality score recalibration were performed using BWA v0.7.12 and GATK v3.3 and following the informed pipeline of the RIG workflow (Auwera et al., 2013; DePristo et al., 2011; Li and Durbin, 2009; McCormick et al., 2015; McKenna et al., 2010). For examining variant accumulation at transcription start sites or coding sequence start sites, the v3.4 gene annotation was used. For all genes, the number of variants at each coordinate relative to the TSS or CDS were summed. For examining periodicity in genome-wide variant accumulation, the average number of variants in a 5,000 bp sliding window centered on the coordinate was determined, then scaled by a factor of 100 (i.e. number of SNPs per 50 base pairs averaged over 5,000 base pairs). To calculate nucleosome occupancy likelihoods, the support vector machine trained by Gupta et al. (2008) was used to calculate likelihoods of 50 bp sliding windows of primary sequence as in Fincher et al. (2013).

Periodicity of SNP accumulation or NOLs was performed using FFTPack within SciPy with the Fast Fourier Transformation (FFT). Genome-wide scans for periodicity were performed using a sliding window of the genome-wide variant accumulation (5,000 bp averages) and NOLs. The signal within a given window was transformed with the FFT, and windows meeting a set of criteria, including strength of a single frequency and a minimum number of cycles, were retained.

### 6.5 Characterization of STA1 and STA2

Sequence corresponding to STA1 and STA2 were identified initially by examining sequence underlying periodic NOLs. The STA1 and STA2 monomers were defined by finding the minimum complete repeat (~180 bp) using BLAST. The starts of the monomers were defined as the region of homology between STA1 and STA2, and for each, the consensus sequence of each monomer was determined by multiple sequence alignment of 9 different monomers representing a trio of tandem repeats from three different arrays on three different chromosome arms (Supplemental Figure S3 and Supplemental File S2) using multalin (Corpet, 1988). Extraction of sequence based on coordinates was facilitated using Biopieces (www.biopieces.org). Internal tandem direct repeats were identified using mreps and YASS (Kolpakov et al., 2003; Noé and Kucherov, 2005).

### 6.6 Gene expression analyses

Gene level read counts were obtained from RNA-seq reads and aligned individually to the version 3 assembly for each biological replicate. The FPKMs of three replicates of a condition were averaged to represent the sample. Per gene FPKMs were analyzed using the scikit-learn python package to perform dimensionality reduction and clustering (Pedregosa et al., 2011). Gene ontology analysis was performed using goatools Python package (Tang et al., 2015).

## 7 DATA ACCESS

The sorghum reference genome sequence and annotation are available from phytozome.jgi.doe.gov. The sequence has also been deposited in GenBank under accession number ABXC00000000. Sequence reads for the 56 resequenced lines are available in the in the National Center for Biotechnology Information Sequence Read Archive (NCBI SRA) under the IDs provided in Supplemental File S5; the 9 lines sequenced as part of this work are associated with BioProject PRJNA374837.

## 1.6 MANUSCRIPT TYPE

Resource

## 8 DISCLOSUE DECLARATION

The authors have no conflicts of interest to declare.

## 9 CONTRIBUTIONS

R.F.M. and S.K.T. performed downstream analyses (e.g. expression clustering, coverage analyses, periodicity analyses), transposon annotation, and linkage analyses. A.S. performed RNA-seq QC, read mapping and expression analyses. S.S. performed gene annotation (gene set version 3.1). J.J, D.S., and J.G. performed genome assembly and finishing (genome version 3.0). M.K. and M.A. performed sorghum transcriptome atlas sequencing. R.F.M., S.K.T., B.W., B.M., and A.M. prepared transcriptome atlas samples. D.M. performed resequencing of selected sorghum lines. J.G., J.S., and J.M. conceived and provided project management. R.F.M., S.K.T., and J.M. wrote the manuscript. All authors reviewed and approved of the manuscript.

## 10 ACKNOWLEDGEMENTS

The work conducted by the U.S. Department of Energy Joint Genome Institute is supported by the Office of Science of the U.S. Department of Energy under Contract No. DE-AC02-05CH11231. This work was funded in part by the DOE Great Lakes Bioenergy Research Center (DOE Office of Science BER DE-FC02-07ER64494), and the U.S. Department of Energy grants no. DE-AR0000596 and DE-SC0012629.

